# Macrophages orchestrate elimination of *Shigella* from the intestinal epithelial cell niche via TLR-induced IL-12 and IFN-γ

**DOI:** 10.1101/2025.01.20.633976

**Authors:** Kevin D. Eislmayr, Charlotte Langner, Fitty L. Liu, Sudyut Yuvaraj, Janet Peace Babirye, Justin L. Roncaioli, Jenna M. Vickery, Gregory M. Barton, Cammie F. Lesser, Russell E. Vance

**Affiliations:** Division of Immunology & Molecular Medicine, Department of Molecular & Cell Biology, University of California, Berkeley, United States; Department of Microbiology, Harvard Medical School, Boston, United States; Broad Institute of Harvard and MIT, Cambridge, United States; Department of Medicine, Division of Infectious Diseases, Massachusetts General Hospital, Boston, United States; Center for Emerging and Neglected Disease, University of California, Berkeley, United States; Cancer Research Laboratory, University of California, Berkeley, United States; Howard Hughes Medical Institute, University of California, Berkeley, United States

**Author notes:** Department of Pathobiology, School of Veterinary Medicine, University of Pennsylvania, Philadelphia, PA, USA.

## Abstract

Bacteria of the genus *Shigella* replicate in intestinal epithelial cells and cause shigellosis, a severe diarrheal disease that resolves spontaneously in most healthy individuals. During shigellosis, neutrophils are abundantly recruited to the gut, and have long been thought to be central to *Shigella* control and pathogenesis. However, how shigellosis resolves remains poorly understood due to the longstanding lack of a tractable and physiological animal model. Here, using our newly developed *Nlrc4*^−/−^*Casp11*^−/−^ mouse model of shigellosis, we unexpectedly find no major role for neutrophils in limiting *Shigella* or in disease pathogenesis. Instead, we uncover an essential role for macrophages in the host control of *Shigella*. Macrophages respond to *Shigella* via TLRs to produce IL-12, which then induces IFN-γ, a cytokine that is essential to control *Shigella* replication in intestinal epithelial cells. Collectively, our findings reshape our understanding of the innate immune response to *Shigella*.

## Introduction

*Shigella* species are the causative agents of shigellosis, a severe inflammatory gastrointestinal infection characterized by symptoms that range from fever, cramps, and nausea to aggressive, mucoid, and bloody diarrhea (dysentery)^1^. Shigellosis is a leading cause of pediatric and geriatric diarrheal-associated mortality, with an estimated 212,000 annual deaths^2–4^. The predominant route of transmission is via the fecal-oral route upon ingestion of contaminated food or water^5^. After initial colonization of the gut, *Shigella* is believed to cross the epithelial barrier through M-cells, a specialized epithelial cell-type sampling the gut lumen. *Shigella* then invades intestinal epithelial cells (IECs) by using its Type III Secretion System (T3SS)^6–8^. This needle-like structure enables the cytosolic delivery of ∼30 virulence effectors that facilitate bacterial entry into and replication within the cytosol of IECs, the preferred intracellular niche for *Shigella*^7,9^. *Shigella* also uses actin-based motility to spread directly from cell-to-cell within the epithelium^10–12^.

Mice are naturally highly resistant to *Shigella* and are able to resist infectious doses 10,000-fold higher than those sufficient to cause disease in humans^13^. Recently, we reported that mice lacking the NAIP–NLRC4 and Caspase-11 inflammasomes are highly susceptible to *Shigella* infection^14,15^. The NAIP–NLRC4 inflammasome detects components of the Shigella T3SS^16–18^, and the Caspase-11 inflammasome senses lipopolysaccharide (LPS) of intracellular *Shigella*^19–22^. Susceptibility to *Shigella* also requires pre-treatment of the inflammasome-deficient mice with antibiotics (e.g., streptomycin) in order to facilitate *Shigella* gut colonization, as is also the case for colonization of mice with other gut pathogens^23–25^. Inflammasomes appear to protect mice from *Shigella* via a highly efficient mechanism in which infected intestinal epithelial cells selectively undergo pyroptotic cell death and expulsion into the gut lumen^26,27^. Although human NAIP and NLRC4 are functional and can detect *Shigella*^28^, the human NAIP–NLRC4 inflammasome does not appear sufficient to protect humans, perhaps due to its poor expression in IECs^29^. Additionally, *Shigella* encodes effectors to block host cell death. For example, the *Shigella* effector IpaH7.8 inactivates human—but not mouse—GSDMD, thus preventing pyroptotic cell death^30^. The *Shigella* effector OspC3 potently blocks CASP4 (the human homolog of mouse Caspase-11)^31^, and partially inhibits mouse Caspase-11^22,31,32^. Mice lacking NAIP–NLRC4 and Caspase-11 (*Nlrc4^−/−^Casp11^−/−^*) thus recapitulate the lack of inflammasome responses in human IECs, and are the only genetically tractable animal model that exhibits all the key manifestations of human shigellosis, including oral infection, efficient bacterial replication in IECs, diarrhea that can be bloody, and neutrophilic inflammation^15^.

*Shigella* infection provokes an acute immune response that involves the secretion of proinflammatory factors such as IL-1β and CXCL1, both of which stimulate the recruitment of myeloid cells^33–35^. As a result, extensive infiltration of neutrophils and monocytes is found in biopsies obtained from humans with active shigellosis, and in infected guinea pigs^36–38^. However, the role of myeloid cells during shigellosis remains poorly understood. Surprisingly, loss of IL-1 signaling did not impact the progression of shigellosis in *Nlrc4*^−/−^ mice^15^. Moreover, *Shigella* largely avoids interacting with phagocytes by replicating within and spreading directly among intestinal epithelial cells. Encounters of *Shigella* with macrophages result in rapid macrophage death^39^. In addition, phagocytosed *Shigella* efficiently escapes phagolysosomal killing by the effector-mediated breakdown of the phagosomal membrane^33,39–42^. For all these reasons, macrophages are not believed to be major participants in bacterial control during shigellosis^7^. By contrast, the massive influx of neutrophils to the gut lumen—one of the primary hallmarks of shigellosis—is believed both to promote bacterial clearance and to cause intestinal damage and severe disease symptoms^7,43^. However, direct experimental assessment of the roles of neutrophils and macrophages during infection is lacking. In particular, a possible role for bystander (uninfected) macrophages has not generally been considered, despite the fact these cells are abundant in the gut and would not be subject to *Shigella*-mediated killing.

Here, we demonstrate that *Shigella* infection in the *Nlrc4^−/−^Casp11^−/−^*mouse model exhibits a self-limiting and spontaneously resolving course of disease very similar to that typically seen in human shigellosis. We unexpectedly discover that in this model neutrophil depletion has no discernable effects on bacterial replication or disease progression, whereas macrophage depletion markedly exacerbates shigellosis symptoms and bacterial burdens in IECs. Our data suggest macrophages orchestrate the innate response to *Shigella* by initiating a protective cytokine circuit. We propose that this circuit begins with bystander (uninfected) macrophages that detect *Shigella* via Toll-like receptors (TLRs), leading to the production of IL-12. IL-12 stimulates the production of IFN-γ, which we demonstrate is an essential factor in resolving the infection *in vivo* and in limiting bacterial replication in both mouse and human intestinal epithelial cells. Mice lacking the ability to produce or sense IFN-γ are incapable of controlling *Shigella* and succumb to the infection. Overall, our study revises our current understanding of *Shigella* pathogenesis and highlights the essential roles of pathogen-sensing by macrophages and the antimicrobial effects of IFN-γ in resolving shigellosis *in vivo*.

## Results

We previously demonstrated that mice deficient in NLRC4 and Caspase-11 (*Nlrc4*^−/−^*Casp11*^−/−^) are susceptible to oral *Shigella* infection and recapitulate the main hallmarks of human shigellosis^14,15^. In humans, *Shigella* infection is typically self-limiting and resolves without intervention within 5-7 days. To assess whether *Shigella* infection of *Nlrc4*^−/−^*Casp11*^−/−^ is also self-limiting, we orally infected streptomycin-pretreated *Nlrc4*^−/−^*Casp11*^−/−^ mice with *Shigella flexneri* strain 2457T and tracked body weight as well as *Shigella* CFU shed in the feces (Figure 1A-B). As expected, after 48h of infection, mice had lost more than 10% of their initial weight and were shedding over 10^9^ *Shigella* CFU/g feces. However, by 72h, mice began to regain weight, coincident with a decrease in *Shigella* CFU in the feces. By day 5, most mice regained or even exceeded their initial weight, and stopped shedding bacteria by day 8 post-infection. These results show that *Nlrc4*^−/−^*Casp11*^−/−^ mice represent a suitable animal model to study both the pathogenesis and resolution of *Shigella* infection.

**Figure 1:**
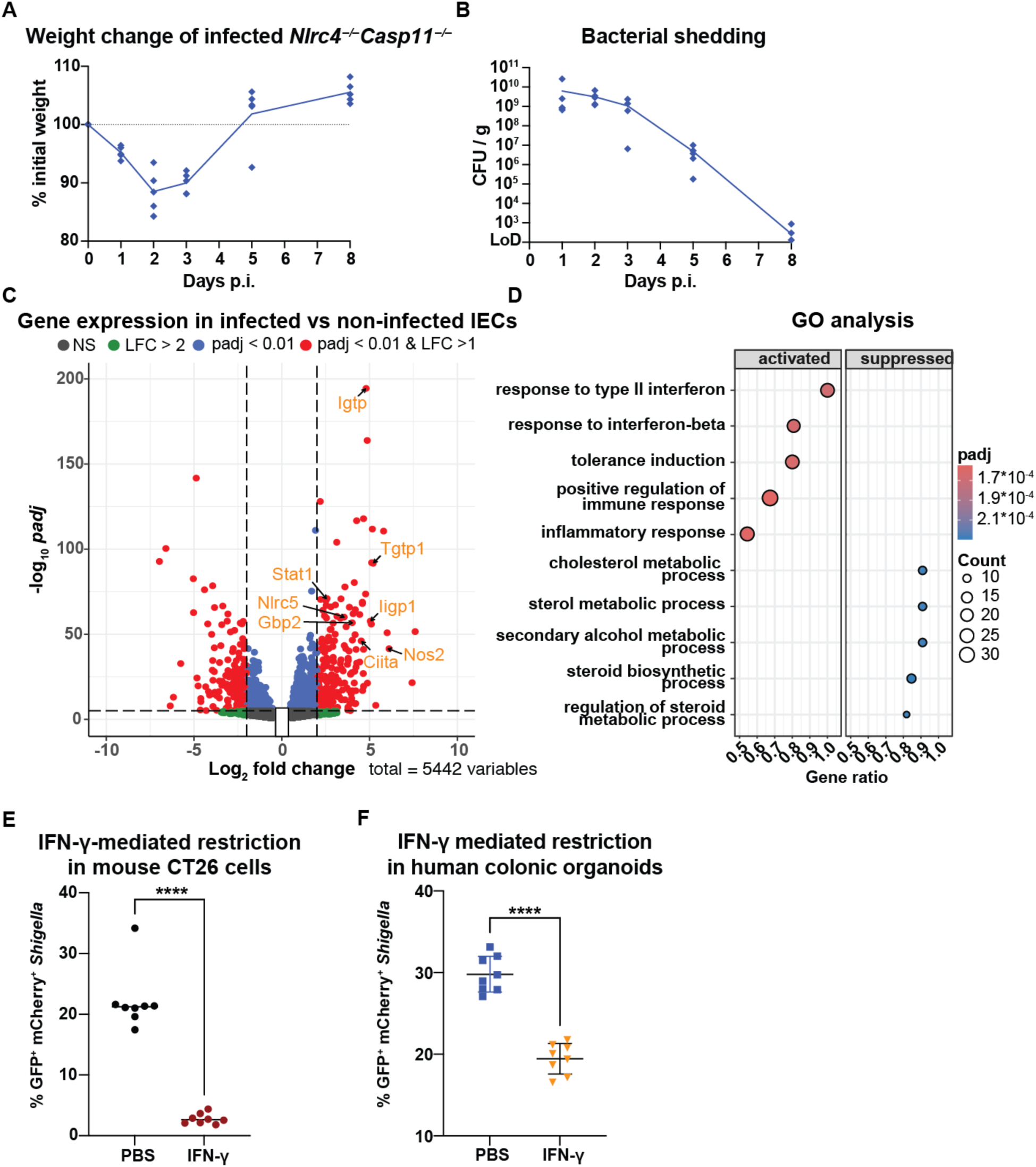
IFN-γ signaling in intestinal epithelial cells is associated with resolution of *Shigella flexneri* infection. **(A)** Body weight (% of the weight on the day of infection) and **(B)** *Shigella* colony forming units (CFU) in the feces of *Nlrc4*^−/−^*Casp11*^−/−^ mice orally infected with *Shigella flexneri*. **(C)** Differential expression analysis of bead-enriched EpCAM^+^ colonic-epithelial cells isolated from infected versus non-infected *Nlrc4*^−/−^*Casp11*^−/−^ mice (n=6 per group, 48h after infection/PBS treatment). Genes with differential expression values with a log_2_ fold change > 1 and padj < 0.001 are highlighted in red. **(D)** Gene ontology (GO) enrichment analysis of significantly differently expressed genes for activated or suppressed biological processes with a gene set size <150. **(E)** *In vitro* infection of mouse CT26 cells with *Shigella* (MOI=1) that constitutively expressing mCherry and an arabinose inducible GFP. Cells were pre-treated with IFN-γ or left untreated for 16 hours. After 45 minutes, gentamycin was added and alive intracellular bacteria were labeled by spiking arabinose into the culture media. The percentage of alive cells containing viable and metabolic active *Shigella* (GFP^+^ mCherry^+^) was assessed by flow cytometry after 3 hours. **(F)** Human colonic organoids were pre-treated with IFN-γ or left untreated for 16 hours and infected with mid-log phase *Shigella* at an MOI of 1. The percentage of mCherry and GFP double-positive cells was assessed by flow cytometry. Median and individual data points in (A) with an n = 5. LoD = Limit of detection. (E, F) with n = 8, ****p<0.0001 in t-test and median shown.

To identify the signaling pathways involved in the restriction of *Shigella* replication, we performed bulk RNA sequencing of bead-enriched EpCAM^+^ epithelial cells isolated from the colon and cecum of mice 48h post-infection. We focused on the cecum and large intestine since bacteria are almost exclusively detected in IECs from these parts of the intestine (Figure S1A). We found 474 genes were significantly upregulated 48h after the infection when compared to naïve mice (log_2_ fold change (lfc) >2, padj < 0.001, Fig. 1C). Among the top differentially expressed genes were *Igtp*, *Tgtp1*, *Nos2* and *Gbp2*, all of which are known to be induced by IFN-γ. Additional interferon-stimulated genes (ISGs) upregulated by *Shigella* infection included *Stat1* and *Ciita*. Indeed, gene set enrichment analysis identified “responses to type II interferon” as the top activated biological process (FDR<0.05, Figure 1D and S1B) for genes with |lfc| > 1, *p*adj <0.05 and a max GO set size <120.

**Figure S1:**
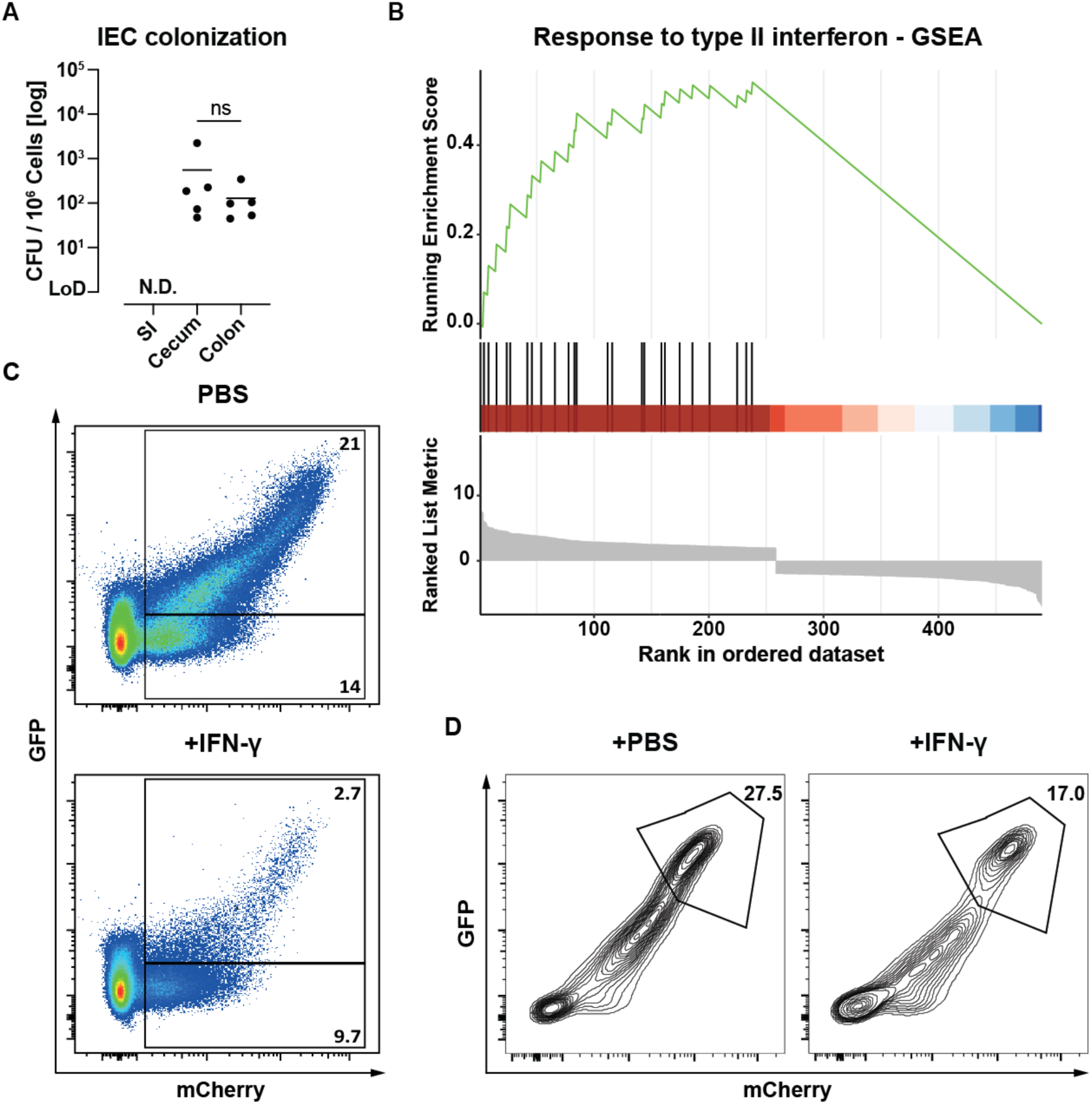
*Shigella* exclusively colonizes the cecum and colon and response to type II interferon limits its intracellular replication *in vitro*. **(A)** Intracellular CFU per million epithelial cells isolated from the small intestine (SI), cecum, and colon, 48 hours after oral infection with *Shigella.* **(B)** Gene set enrichment analysis (GSEA) for the biological process “Responses to type II interferon” identified in Fig. 1D plotted as running enrichment scores (ES) and positions of gene set members on the rank-ordered list of differently expressed genes in IECs isolated from *Shigella* infected and naïve mice. **(C**) Representative flow cytometry plots for quantifying infected (GFP^+^mCherry^+^) CT26 cells in Fig. 1E with % of cells within the corresponding gate**. (D)** Representative flow cytometry plots for quantifying infected (GFP^+^mCherry^+^) human colonic organoids in Fig. 1F with. (A) n = 5, Mann-Whitney-test with the line indicating the median. LoD = Limit of detection, N.D. = no CFUs detected.

### IFN-γ restricts *Shigella* replication *in vivo* and protects against shigellosis disease

Consistent with previous results demonstrating that IFN-γ inhibits the growth of *S. flexneri* in mouse embryonic fibroblasts^44^, we found that pretreatment of CT26 cells (a mouse colon carcinoma cell line) with IFN-γ for 16 hours prior to infection resulted in substantially fewer cells harboring metabolically active *Shigella* 3 hours after infection (Fig. 1E, Fig. S1C). Similarly, 16-hour pretreatment of human colonic organoids with IFN-γ significantly reduced the proportion of *Shigella*-infected cells (Fig. 1F, Fig. S1D).

*In vivo*, we observed that IFN-γ levels in the lamina propria (LP) were detectable as early as 8 hours after infection, and continued to rise throughout the infection (Fig. 2A), suggesting IFN-γ may act early to restrict *Shigella* replication. To test whether IFN-γ is important to resolve *Shigella* infection, we treated *Nlrc4*^−/−^*Casp11*^−/−^ mice with an IFN-γ-neutralizing antibody every 24 hours throughout the infection (starting at day 0). Anti-IFN-γ-treated mice lost significantly more weight than control animals injected with an isotype control antibody (Fig. 2B), and their IECs harbored >100-fold more *Shigella* CFU (Fig. 2C). Colonic and cecal atrophy—a hallmark of intestinal inflammation—was more pronounced in IFN-γ-neutralized mice (Fig. 2D, S2A). We also noticed that 5 out of 8 animals treated with anti-IFN-γ exhibited high levels of occult blood (strong positive test, score = 2; see Methods for scoring scale) and the remaining 3 were tested positive for occult blood (faintly positive test, score = 1) (Fig. 2E). In comparison, only 50% of control animals exhibited a positive occult blood test, and these were only faintly positive (score of 1). Of note, mice treated with anti-IFN-γ harbored much higher levels of live bacteria in their mesenteric lymph nodes and spleens (Fig. 2F). Moreover, fecal levels of calprotectin, lipocalin and myeloperoxidase (MPO), indicative of intestinal inflammation, were elevated in IFN-γ-neutralized mice (Fig. 2G), as were the levels of inflammatory cytokines (IL-1β and CXCL1) in the lamina propria (Fig. 2H). Notably, *Shigella* CFU within the feces remained similar between anti-IFN-γ and isotype-treated mice, indicating that altered intestinal luminal colonization did not explain the effects of IFN-γ neutralization (Fig. 2I). From these results, we conclude that IFN-γ is critical to restrict intracellular *Shigella* replication and prevent severe disease in *Nlrc4*^−/−^*Casp11*^−/−^ mice *in vivo*.

**Fig. 2:**
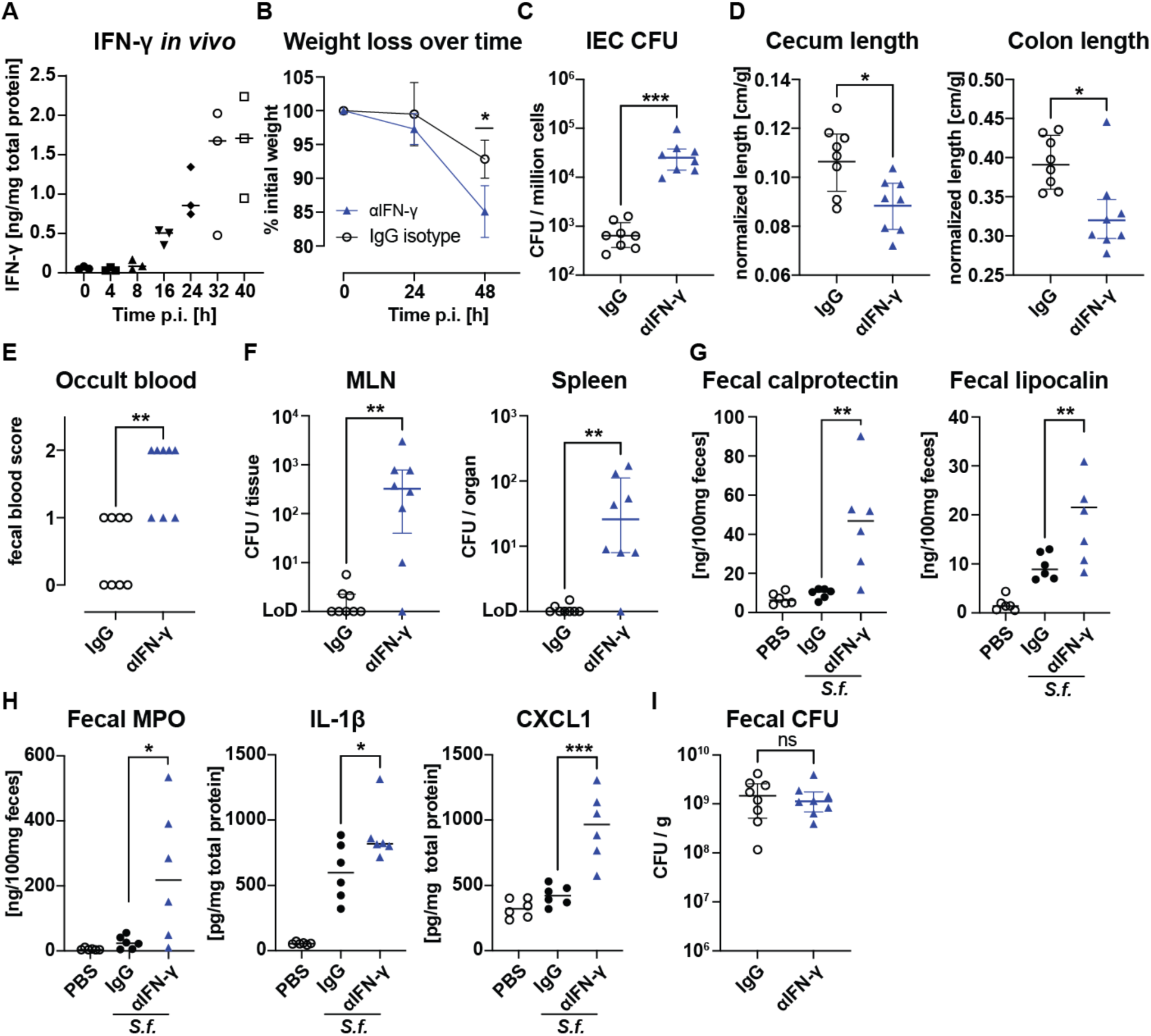
IFN-γ is essential to protect from severe disease and limit bacterial replication during shigellosis. **(A)** ELISA quantification of IFN-γ levels in the lamina propria of infected *Nlrc4*^−/−^*Casp11*^−/−^ mice at different time points post-infection (n = 3 per time point). Mice were treated with anti-IFN-γ antibody or isotype control IgG (500µg injected i.p. every 24h starting at 0h.p.i.) and were assessed for **(B)** body weight, **(C)** intracellular bacteria in cecum/colon epithelial cells, **(D)** length of cecum and colon normalized to the initial weight of infected mice treated with IFN-γ neutralizing antibody or an IgG control, **(E)** fecal occult blood score (see methods for scale), **(F)** bacterial dissemination into the mesenteric lymph nodes (MLN) and the spleen, **(G, H)** clinical markers of gut inflammation (calprotectin, lipocalin and MPO in the feces) as well as the pro-inflammatory cytokines IL-1β and CXCL1 levels in lamina propria, measured 48 hours after infection treated with IFN-γ neutralizing antibody or isotype control. PBS-treated non-infected mice were added to assess cytokine levels at steady state. **(I)** Quantification of shed Shigella bacteria in the fecal matter of infected animals. n = 6 PBS and 8 infected mice per treatment group. *p < 0.05, **p < 0.01, ***p < 0.001 in t-test with mean and SD shown (B,D,G,H) or Mann-Whitney-test with median and interquartile range (C,E,F,I). ns = not significant (p>0.05).

**Figure S2:**
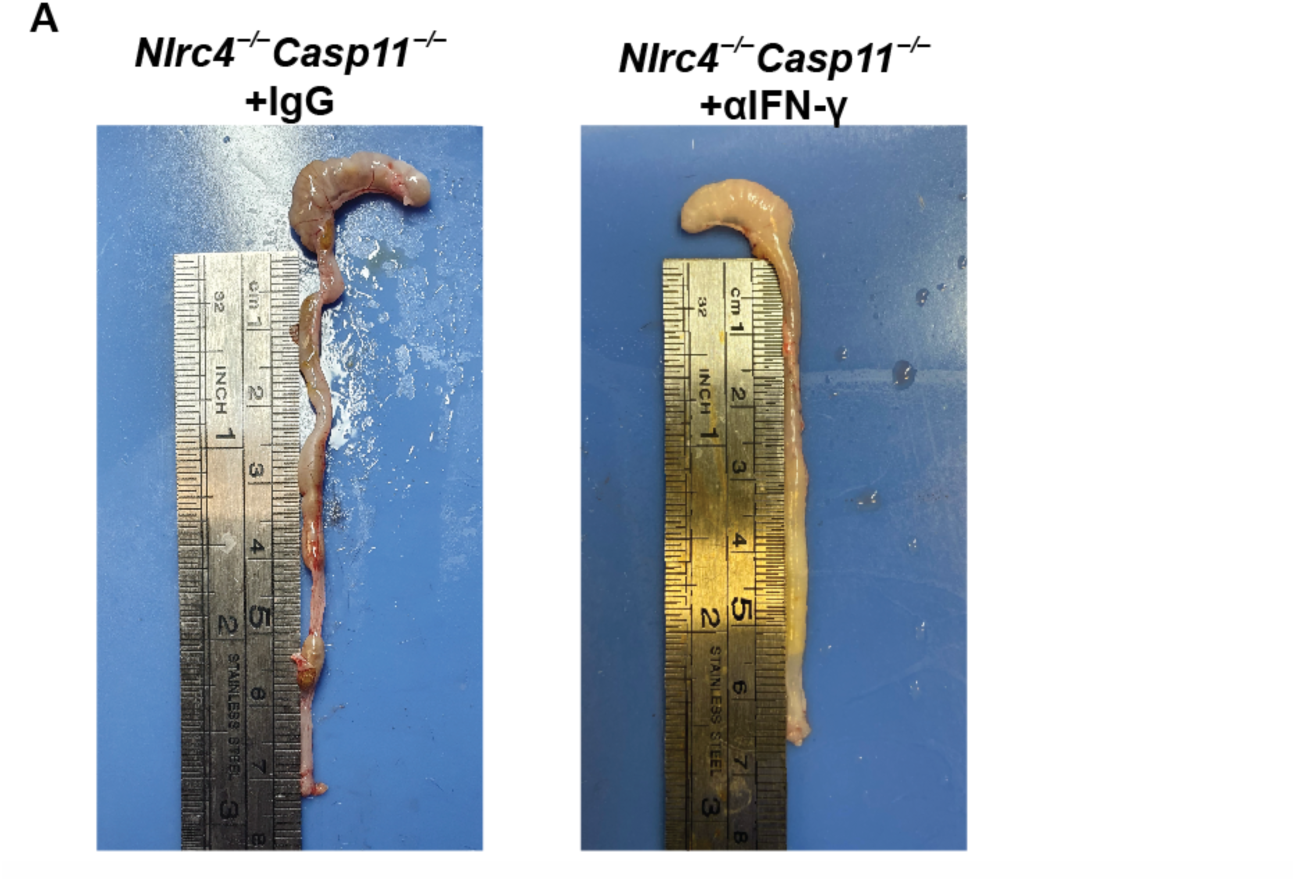
IFN-γ neutralization enhances atrophy. Representative images of cecum and colon isolated from *Nlrc4*^−/−^*Casp11*^−/−^ mice 48 hours after oral infection with *Shigella* and treatment with IFN-γ-neutralizing antibody (αIFN-γ, right) or matching isotype control (IgG, left)

### IFN-γ is essential for *Shigella* control and acts predominantly on non-hematopoietic cells

To provide genetic evidence that IFN-γ is essential for *Shigella* control, we generated *Nlrc4*^−/−^ *Casp11*^−/−^ mice that also lack the IFN-γ-receptor (*Ifngr1*). Consistent with the effects of IFN-γ neutralization (Fig. 2), we found that IECs from *Shigella*-infected *Ifngr1*^−/−^*Nlrc4*^−/−^*Casp11*^−/−^ mice harbor significantly more bacteria than IFN-γ-sufficient *Nlrc4*^−/−^*Casp11*^−/−^ animals 48 hours after infection. Treatment of *Ifngr1*^−/−^*Nlrc4*^−/−^*Casp11*^−/−^ mice with anti-IFN-γ did not further enhance susceptibility, suggesting that anti-IFN-γ does not have non-specific effects (Fig. S3A). *Ifngr1*^−/−^ *Nlrc4*^−/−^*Casp11*^−/−^ mice experienced significantly greater weight loss at 48 hours post-infection than *Nlrc4*^−/−^*Casp11*^−/−^ mice (Fig. 3A). Strikingly, by 72 hours post-infection, all *Ifngr1*^−/−^*Nlrc4*^−/−^ *Casp11*^−/−^ mice reached predefined humane endpoints (e.g., <75% starting body weight, severe signs of morbidity) and had to be euthanized, whereas all *Nlrc4*^−/−^*Casp11*^−/−^ mice survived the infection and regained their initial weight by 120 hours after infection (Fig. 3B). Together, our data indicate a critical role for IFN-γ in restricting *Shigella* replication and in promoting recovery from shigellosis.

**Figure 3.**
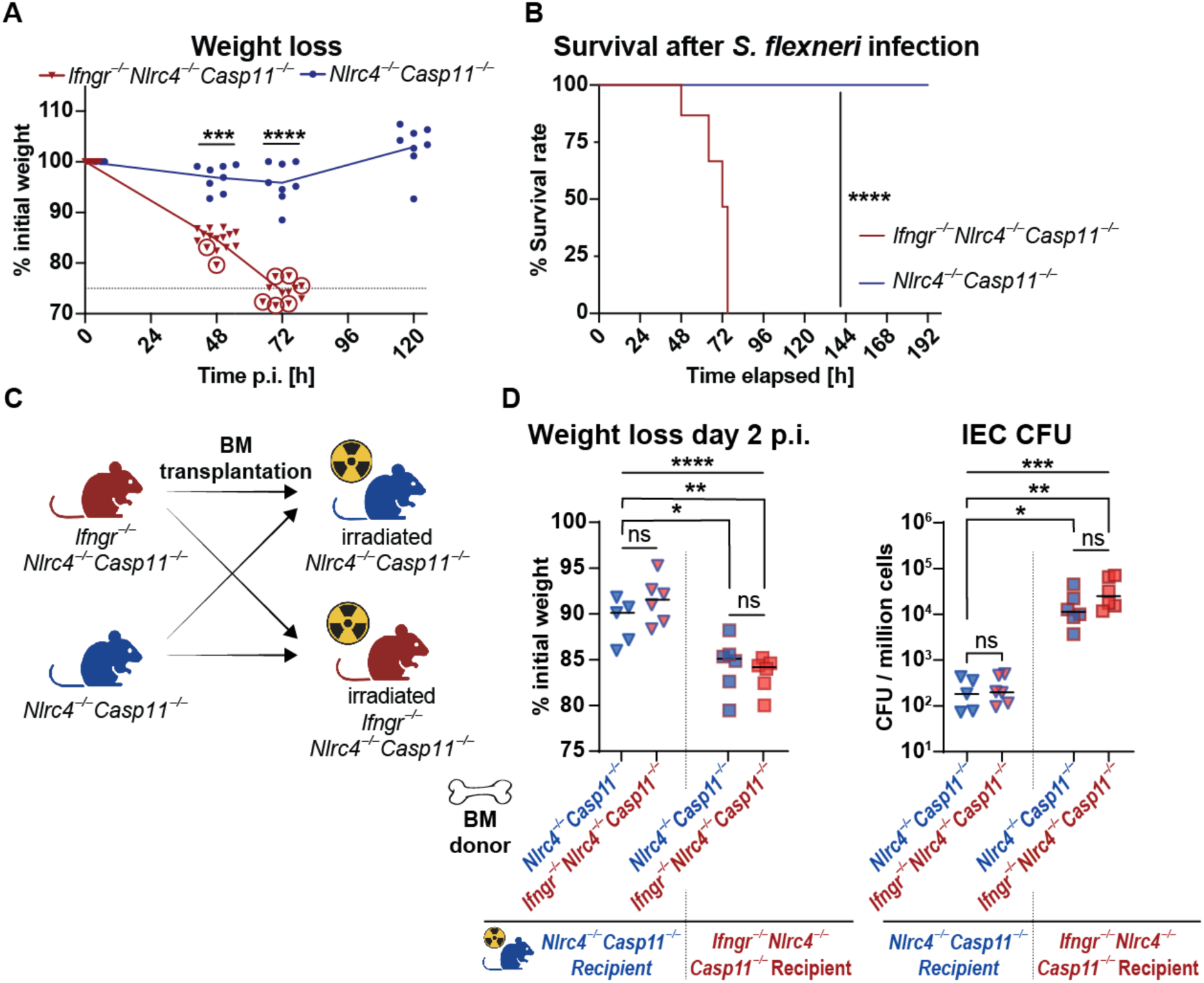
Responsiveness of radioresistant (non-hematopoietic) cells to IFN-γ is essential for the control of *Shigella* infection. Resistance to *Shigella* infection of *Nlrc4*^−/−^*Casp11*^−/−^ mice compared to animals additionally lacking the IFN-γ receptor (*Ifngr1*^−/−^*Nlrc4*^−/−^*Casp11*^−/−^) was assessed by **(A)** the percentage of mice exceeding human endpoint criteria (indicated with circled symbols) and **(B)** body weight. The dashed line represents the predefined 75% weight retention limit set as a humane endpoint. **(C)** Experimental design of reciprocal bone marrow transplantation, **(D)** the change in weight and number of colonic intra-epithelial CFU 48 hours after infection. The left half of each graph represents data obtained from irradiated *Nlrc4*^−/−^*Casp11*^−/−^ mice engrafted with either *Nlrc4*^−/−^ *Casp11*^−/−^ or *Ifngr1*^−/−^*Nlrc4*^−/−^*Casp11*^−/−^ bone marrow, whereas the right portion is from irradiated *Ifngr1*^−/−^*Nlrc4*^−/−^*Casp11*^−/−^ mice engrafted with *Nlrc4*^−/−^*Casp11*^−/−^ or *Ifngr1*^−/−^*Nlrc4*^−/−^*Casp11*^−/−^ bone marrow, respectively. n = 15 *Ifngr1*^−/−^*Nlrc4*^−/−^*Casp11*^−/−^ and 8 *Nlrc4*^−/−^*Casp11*^−/−^ in (A, B). ***p<0.001, ****p<0.0001 in t-test (A) and Log-rank (Mantel-Cox) test (B). Data from 5-6 mice per group in (D) with mean and one-way ANOVA with Tukey’s multiple comparison for weight and median and Kruskal-Wallis test with Dunn’s multiple comparison for CFU. *p<0.05, **p<0.01, ns = not significant (p>0.05)

Next, we sought to determine whether IFN-γ acts on hematopoietic or non-hematopoietic cells to restrict *Shigella* replication *in vivo*. We therefore generated reciprocal bone marrow chimeras between *Ifngr1*^−/−^*Nlrc4*^−/−^*Casp11*^−/−^ mice and *Nlrc4*^−/−^*Casp11*^−/−^ mice (schematic in Fig. 3C). After allowing 8 weeks for the hematopoietic compartment to reconstitute, we infected the chimeric mice with *Shigella*. Examining the mice 48 hours after infection revealed that IFN-γ-mediated resistance to *Shigella* depended on the genotype of the bone marrow recipient, regardless of the genotype of the donor (Fig. 3D). Recipients lacking the IFN-γR phenocopied full-body IFN-γR deficiency, regardless of whether they were reconstituted with *Ifngr1*^−/−^*Nlrc4*^−/−^ *Casp11*^−/−^ or *Nlrc4*^−/−^*Casp11*^−/−^ bone marrow. Conversely, *Shigella* replication in IECs was restricted in irradiated *Nlrc4*^−/−^*Casp11*^−/−^ mice reconstituted with either *Ifngr1^−/−^Nlrc4*^−/−^*Casp11*^−/−^ or *Nlrc4*^−/−^*Casp11*^−/−^ bone marrow. From these results, we conclude that IFN-γ predominantly acts on radioresistant (likely non-hematopoietic) cells to restrict *Shigella* replication. Given that we observe profound upregulation of ISGs in intestinal epithelial cells during infection (Fig. 1C), our data are consistent with the hypothesis that IFN-γ restricts *Shigella* replication by eliciting an anti-bacterial response in IECs.

**Figure S3:**
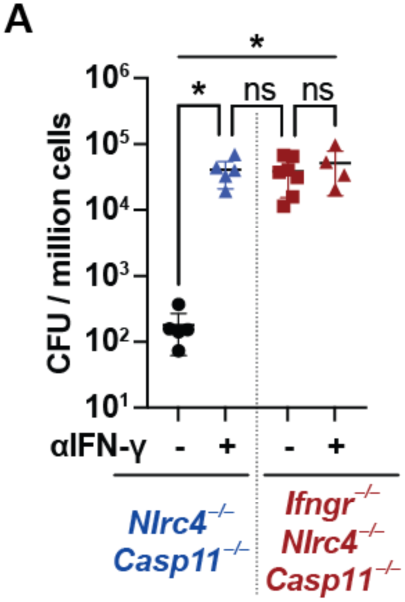
IFN-γ neutralizing antibody has no effect in the absence of the IFN-γ receptor (IFNGR1) **(A)** Intracellular bacterial counts in IECs from infected *Ifngr1*^−/−^*Nlrc4*^−/−^*Casp11*^−/−^ or *Nlrc4*^−/−^ *Casp11*^−/−^ mice treated or not treated with IFN-γ neutralizing antibody. n = 4-7 mice pre-group, Kruskal-Wallis test with Dunn’s multiple comparison. *p<0.05, ns = not significant (p>0.05)

### Macrophages are essential for *Shigella* control, but neutrophils are dispensable

Although hematopoietic cells did not appear to be critical responders to IFN-γ, we hypothesized that immune cells may still participate in the signaling circuit leading to the production of IFN-γ. We observed that large numbers of neutrophils (Ly6G^+^Ly6C^med^) and inflammatory monocytes (Ly6G^−^Ly6C^high^MHCII^+^) migrate into the lamina propria after *Shigella* infection (Fig. S4A). The number of neutrophils increased by 15-fold, and inflammatory monocytes by 5-fold, compared to naïve mice, while the number of T-cells remained similar.

To determine if myeloid cells contribute to the clearance of *Shigella*, we generated *Nlrc4*^−/−^ *Casp11*^−/−^ mice in which a Cre-inducible diphtheria toxin receptor gene is specifically expressed by LysM^+^ (myeloid) cells (*Lyz2^Cre/+^iDTR^lsl/lsl^Nlrc4*^−/−^*Casp11*^−/−^ mice). Treatment of these mice with diphtheria toxin (DT) results in the depletion of myeloid cells, primarily macrophages and neutrophils (Figure S4B, S4C). *Shigella* infection of LysM^+^ myeloid cell-depleted mice resulted in a significantly higher weight loss and ∼100-fold more *Shigella* CFUs in IECs, as compared to non-depleted mice (Fig. 4A). In addition, LysM^+^ myeloid cell depletion resulted in increased inflammatory cytokines and markers, indicating an overall exacerbation of the pathology (Figure S4D). LysM^+^ myeloid cell depletion also resulted in significantly enhanced levels of occult/visible blood in the feces (Fig. 4B). Thus, we conclude that myeloid cells critically contribute to the control of shigellosis.

**Figure 4.**
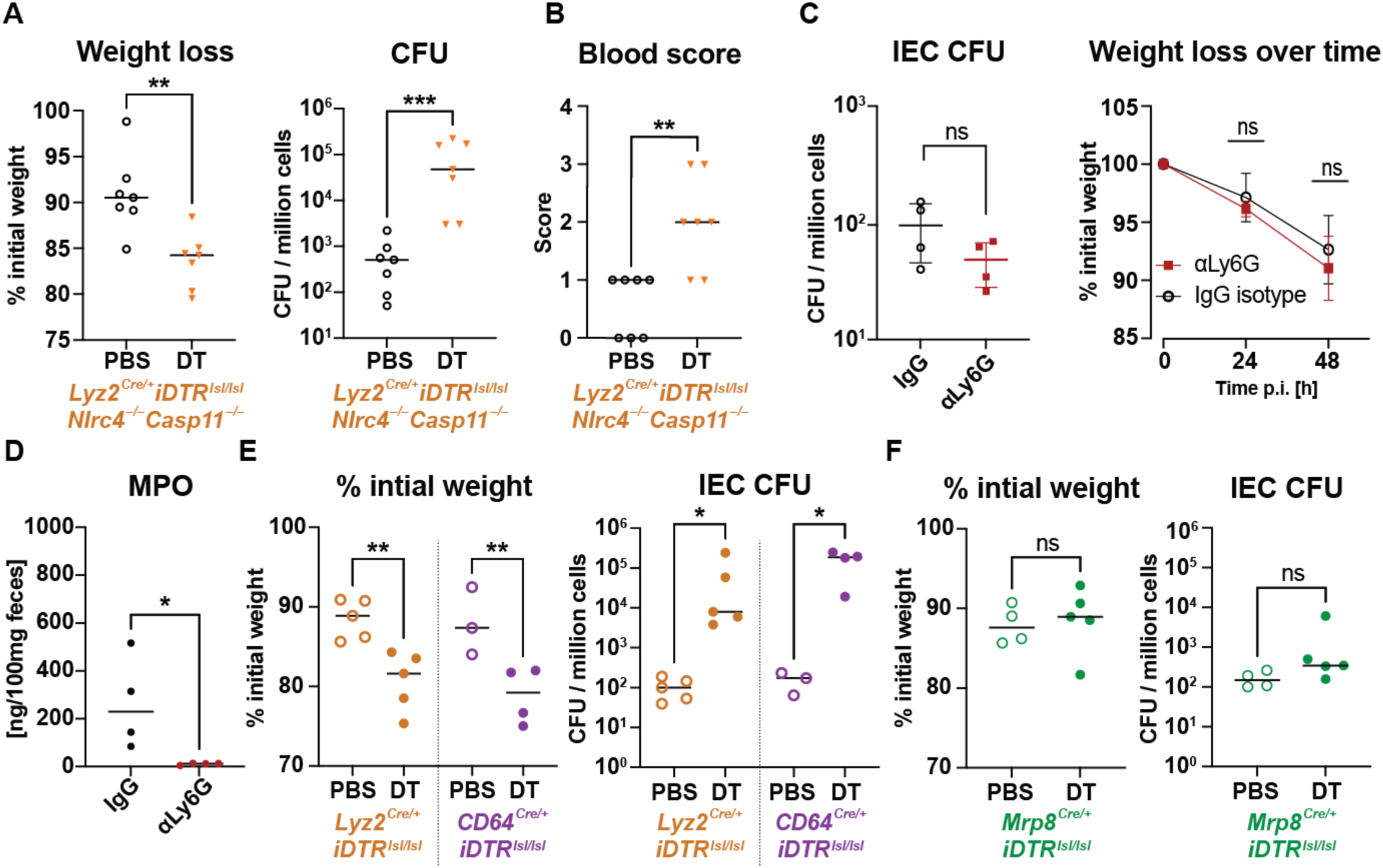
LysM^+^/CD64^+^ myeloid cells, but not neutrophils, are essential for the control of shigellosis. **(A)** Change in body weight and epithelial intracellular bacterial counts and **(B)** fecal occult blood score of infected *Lyz2^Cre/+^iDTR^lsl/lsl^Nlrc4*^−/−^*Casp11*^−/−^ mice in myeloid cell-depleted (DT-treated; orange triangle) or non-depleted (without DT treatment; open black circle) mice. Blood score is based on the result of a fecal occult blood test (stratified from 0=negative, 1=slightly positive, 2= strongly positive test result) or the presence of visual blood (score = 3) in the lumen of the large intestine. **(C)** IEC CFU counts 48 hours after infection and weight over time of *Shigella*-infected *Nlrc4*^−/−^*Casp11*^−/−^ mice treated with a neutrophil depleting antibody (αLy6G) or an IgG isotype control. **(D)** Effect of neutrophil depletion on fecal MPO levels 48 hours after infection. **(E)** Body weight and bacterial colonization of epithelial cells of irradiated *Nlrc4*^−/−^*Casp11*^−/−^ mice engrafted with *Lyz2^Cre/+^iDTR^lsl/lsl^*, or *CD64^Cre/+^iDTR^lsl/lsl^*bone marrow, respectively, treated with DT or PBS (n = 4 per group). **(F)** Weight loss and IEC CFU counts obtained 48 hours after infection from irradiated *Nlrc4*^−/−^*Casp11*^−/−^ mice reconstituted with *Mrp8^Cre/+^iDTR^lsl/lsl^* bone marrow. Mice were treated with DT to specifically deplete neutrophils, or with PBS as a negative control. Data from experiments with n = 7 (A-B), n = 4 (C-D), n = 3-5 (E), n = 4-5 (F). *p<0.05, **p<0.01, ***p<0.001, ns = not significant according to t-test (A,D, weight data in F) with mean and SD sown or Mann-Whitney-test (CFU data in A,B,C,F) with median and interquartile range, one-way ANOVA with Tukey’s multiple comparisons for weight (E) and median and Kruskal-Wallis test with Dunn’s multiple comparison for CFU (E).

Neutrophils are believed to play important roles in controlling bacterial burdens and in causing pathology during shigellosis. Moreover, we observe significant recruitment of neutrophils to the gut tissue of our *Shigella*-infected *Nlrc4*^−/−^*Casp11*^−/−^ mice (Figure S4A). Therefore, we assessed the impact of neutrophil depletion during infection using a Ly6G-specific antibody (clone 1A8). Unexpectedly, we observed that neutrophil depletion did not affect bacterial burdens within IECs compared to non-treated animals (Fig. 4C). Furthermore, neutrophil depletion did not impact weight loss or the induction of inflammatory markers during *Shigella* infection (Fig. 4C and Fig. S4E). We confirmed that neutrophils were depleted by 1A8 treatment (Fig. S4F). Consistent with successful neutrophil depletion, 1A8-treated mice did exhibit reduced pus and MPO levels in the cecum and colon (Fig. 4D, representative images in Fig. S4H). However, neutrophil depletion did not affect the levels of *Shigella* in the gut lumen (Fig. S4G).

To verify the results of antibody depletion, we crossed the neutrophil-specific *MRP8-Cre-ires-GFP* (*Mrp8^Cre/+^*) animals^45^ with *iDTR^lsl/lsl^* mice to generate *Mrp8^Cre/+^iDTR^lsl/lsl^* mice. Irradiated *Nlrc4*^−/−^*Casp11*^−/−^ mice were then engrafted with *Mrp8^Cre/+^iDTR^lsl/lsl^* bone marrow. After reconstitution, the chimeric mice were infected with *Shigella* and treated with DT to deplete *Mrp8*^+^ cells (neutrophils). Surprisingly, DT treatment of infected mice did not alter weight loss or IEC CFU burdens as compared to PBS controls (Fig. 4F). DT treatment efficiently depleted neutrophils in the blood and lamina propria of infected mice (Fig. S4I, J). Thus, our results indicate that neutrophil depletion has no major effect on the pathogenesis of *Shigella* infection.

Based on the significant impact on IEC colonization observed with the *Lyz2^Cre/+^iDTR^lsl/lsl^* model and the apparently negligible role of neutrophils, we proceeded to investigate the role of macrophages and monocytes in controlling *Shigella* infection. Scott *et al.* recently described the use of CD64^Cre^ mice for macrophage-specific Cre expression^46^. Therefore, we crossed *CD64^Cre/+^* mice to *iDTR^lsl/lsl^*animals and subsequently transplanted the bone marrow into irradiated *Nlrc4*^−/−^ *Casp11*^−/−^ mice (*CD64^Cre/+^iDTR^lsl/lsl^* into *Nlrc4*^−/−^*Casp11*^−/−^ chimeras). DT-treated (CD64^+^ cell depleted) infected mice experienced more pronounced weight loss compared to PBS-treated (non-depleted) mice (Fig. 4E). Moreover, DT-mediated CD64^+^ cell depletion led to an IEC CFU burden similar to that seen with LysM^+^ cell depletion. Notably, the lamina propria of infected and DT-injected *CD64^Cre/+^iDTR^lsl/lsl^* chimeras exhibited a significant absence of neutrophils in addition to the depletion of CD64^+^ macrophages, in both the LAP and blood (Fig. S4I, J), whereas no effect of DT treatment on blood neutrophils was seen in naïve mice (Fig. S4I, J). Hence, although CD64^Cre^ is specific for monocytes and macrophages under non-inflamed conditions^46^, it may be expressed by neutrophils during inflammation. However, in our experiments, we rule out an essential role for neutrophils with neutrophil-specific depletion by 1A8 and *Mrp8^Cre/+^iDTR^lsl/lsl^*mice. We conclude that macrophages play a central role in resistance to *Shigella* infection.

**Figure S4:**
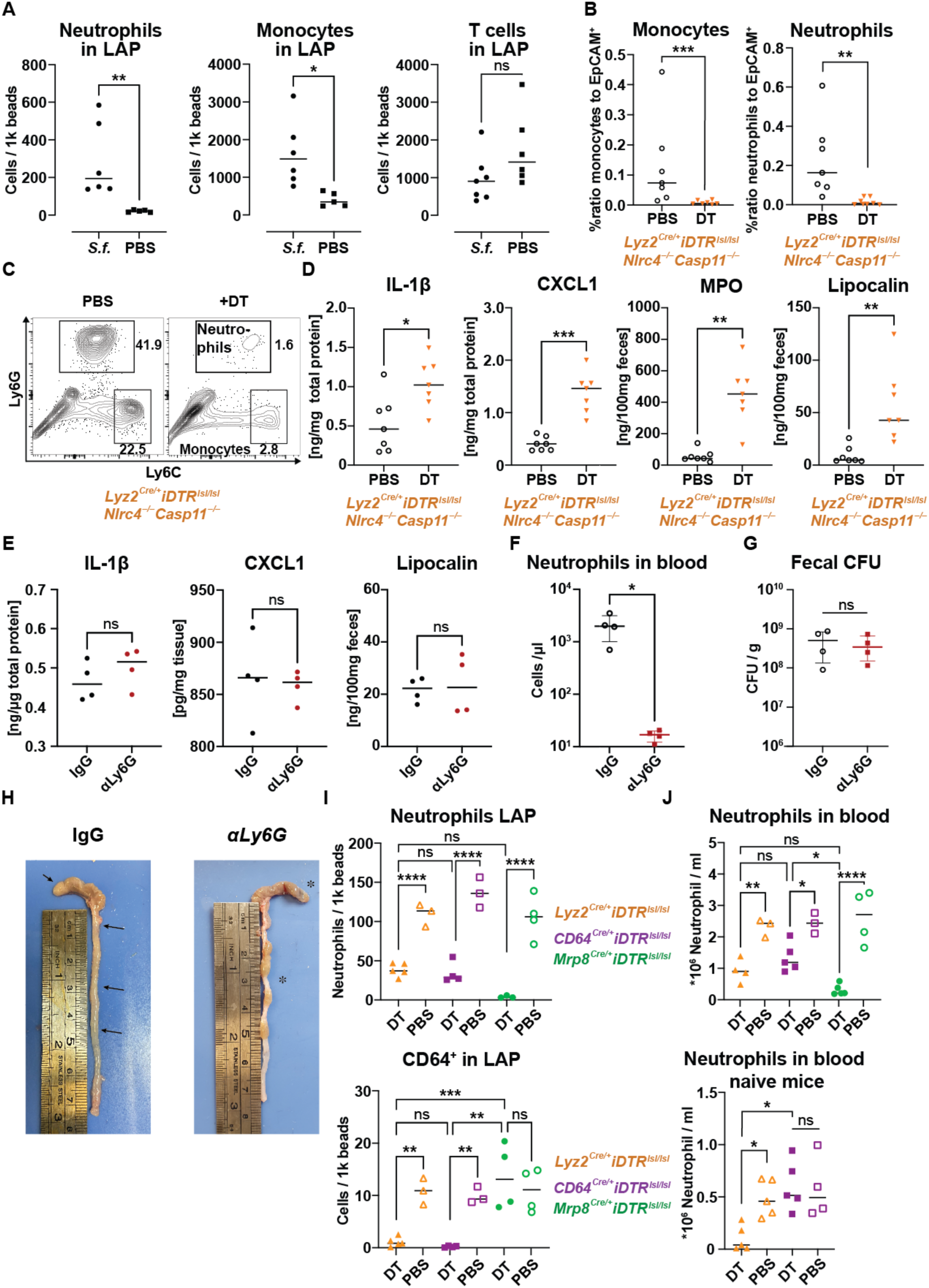
Increase bowel inflammation in myeloid-deficient mice. **(A)** Neutrophils, monocytes and T cells in the lamina propria (LAP) of infected (*S.f.*) or naïve (PBS) *Nlrc4*^−/−^*Casp11*^−/−^ mice. At the time of infection, mice were treated with an IFN-γ-neutralizing antibody or an isotype control. **(B)** Monocyte and neutrophil levels in the colon/cecum of infected *Lyz2^Cre/+^iDTR^lsl/lsl^Nlrc4*^−/−^*Casp11*^−/−^ mice with or without diphtheria toxin treatment **(C)** representative flow plots of the data in B. **(D)** Effect of myeloid cell depletion on IL-1b, CXCL1 protein levels in the LAP, and MPO and Lipocalin in the feces. **(E)** Impact of neutrophil depletion (anti-Ly6G clone 1A8) on levels of IL-1β and CXCL1 in the LAP, and lipocalin in the feces, of infected *Nlrc4*^−/−^*Casp11*^−/−^ mice. **(F)** Quantification of circulating neutrophils in the blood and **(G)** luminal colonization of infected animals treated with anti-Ly6G or an isotype control. **(H)** Representative images of cecum and colon of *Nlrc4*^−/−^*Casp11*^−/−^ mice treated with anti-Ly6G or an isotype control 48 hours post-infection. Arrows indicate sticky, white-to-yellow pus observed in infected *Nlrc4*^−/−^*Casp11*^−/−^ (left) that is absent upon antibody-mediated neutrophil depletion (right). **(I)** Ly6G^+^ neutrophils (top) and CD64^+^ macrophages in the LAP from infected irradiated *Nlrc4*^−/−^*Casp11*^−/−^ mice, engrafted with *Lyz2^Cre/^ iDTR^lsl/lsl^*, *CD64^Cre/+^iDTR^lsl/lsl^*, or *MRP8^Cre/+^iDTR^lsl/lsl^* bone marrow, and treated with DT or PBS (n=3-4 per group). **(J)** Ly6G^+^ neutrophils in the blood of irradiated *Nlrc4*^−/−^*Casp11*^−/−^ mice, engrafted with *Lyz2^Cre/+^iDTR^lsl/lsl^*, *CD64^Cre/+^ iDTR^lsl/lsl^*, or *MRP8^Cre/+^ iDTR^lsl/lsl^* bone marrow, and treated with DT or PBS, 48h after infection with *Shigella* (top) or before infection (bottom). n=5-6 in A, n=7 per group in B and D, n=4 in E-G, and n=3-5 in I-J. *p<0.05, **p<0.01, ***p<0.001, ns=not significant in Mann-Whitney-test for (A,B,D,F,G) with median shown, t-test for (E) and ANOVA with Dunnett’s multiple comparisons test in (I,J).

### IFN-γ production is dependent on macrophage TLR sensing

Since both IFN-γ neutralization and depletion of macrophages/monocytes led to heightened bacterial IEC colonization, we wondered whether macrophages/monocytes might play a role in IFN-γ induction during infection. Indeed, IFN-γ levels were significantly reduced in the LAP of myeloid cell-depleted *Lyz2^Cre/+^iDTR^lsl/lsl^Nlrc4*^−/−^*Casp11*^−/−^ mice (Fig. 5A). These data suggest that the induction of IFN-γ is a major contribution of macrophages and monocytes in restricting *Shigella*.

**Figure 5:**
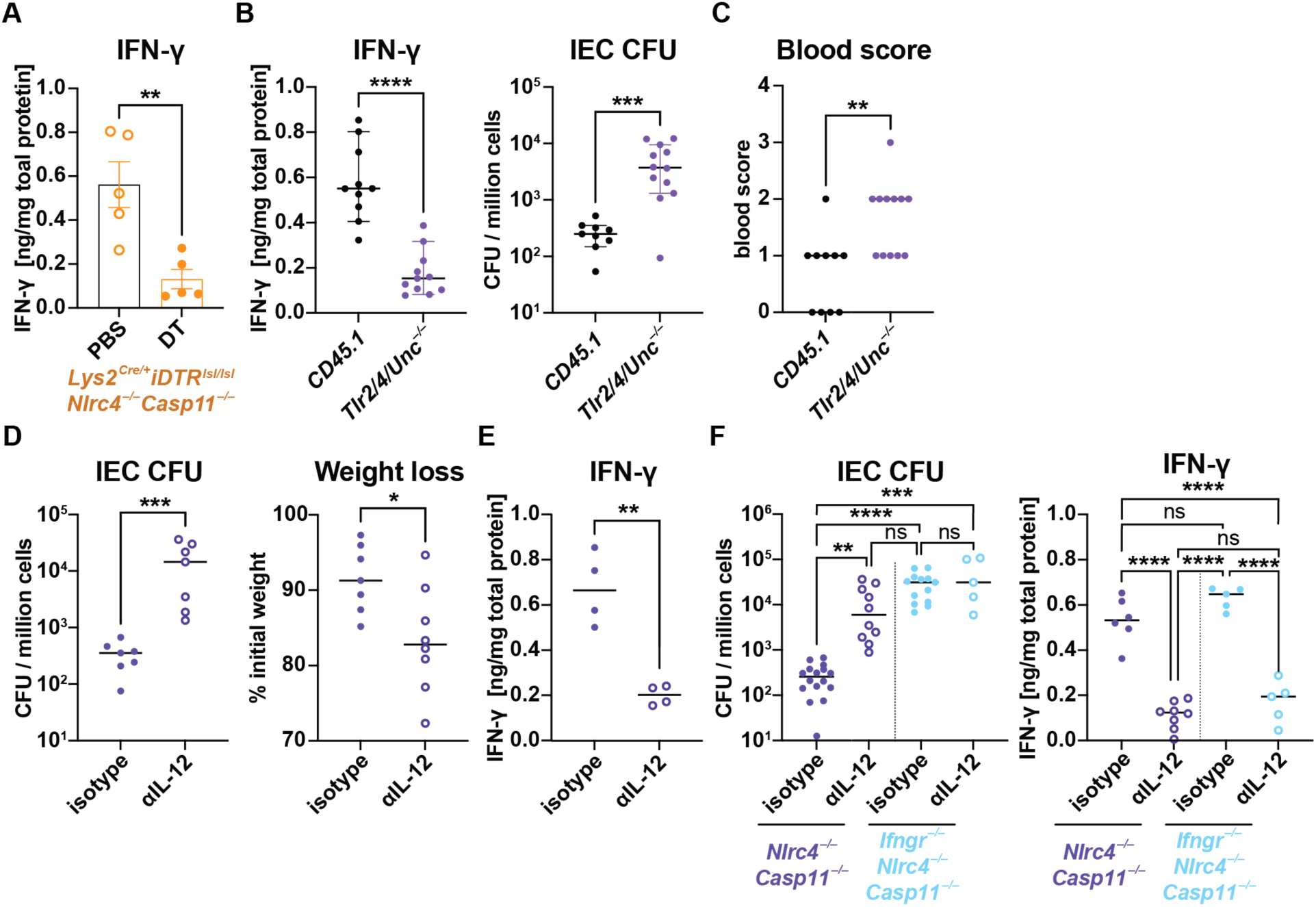
TLR and IL-12 signaling are important for IFN-γ production and bacterial control during *Shigella* infection. **(A)** IFN-γ levels in the lamina propria of infected *Lyz2^Cre/+^iDTR^lsl/lsl^Nlrc4*^−/−^*Casp11*^−/−^ mice with or without DT-induced myeloid cell depletion. **(B)** IFN-γ in the LAP and IEC CFU 48 hours post-infection in irradiated *Nlrc4*^−/−^*Casp11*^−/−^, receiving *Tlr2/4-Unc^−/−^* or CD45.1-WT bone marrow and **(C)** the corresponding fecal occult blood score. **(D)** Effect of IL-12 neutralization (αIL-12) or isotype treatment during *Shigella* infection of *Nlrc4*^−/−^*Casp11*^−/−^ mice on IEC CFU counts, weight loss and **(E)** IFN-γ levels in LAP 48 hours after infection. **(F)** Comparison of αIL-12 or isotype treatment in *Nlrc4*^−/−^*Casp11*^−/−^ or *Ifngr^−/−^Nlrc4*^−/−^*Casp11*^−/−^ mice regarding the observed IEC CFU counts and IFN-γ levels in the LAP. *p<0.05, **p<0.01, ***p<0.001, ****p<0.0001, ns=not significant according to t-test (A,F), Mann-Whitney-test (B CFU, C, E CFU and weight loss), Dunn’s multiple comparisons test (CFU data in G) and Tukey’s multiple comparisons test (IFN-γ data in G). n=5 (A), n=9-12 (B), n=7 (E,F), n=5-16 from two independent experiments (G).

Previous studies have established that macrophages sense bacterial infection via Toll-like receptors (TLRs) and produce pro-inflammatory cytokines (including the IFN-γ-inducing cytokine, IL-12). Thus, we addressed the importance of TLR signaling for IFN-γ production. It has been demonstrated that *Unc93b1*-deficient cells no longer respond to TLR3, TLR7, TLR9, TLR11, TLR12, and TLR13 ligands^47^. By crossing these mice to *Tlr2*^−/−^ and *Tlr4*^−/−^ mice, Sivick et al generated mice incapable of TLR signaling^48^, while still retaining the capacity for IL-1R and IL-18R signaling (which, like TLRs, signal via MyD88). *Tlr2/4/Unc*^−/−^ into *Nlrc4*^−/−^*Casp11*^−/−^ bone marrow chimeras displayed blunted IFN-γ induction compared to WT(CD45.1) into *Nlrc4*^−/−^ *Casp11*^−/−^ control chimeras (Fig. 5B). Consistent with the important role of IFN-γ in the control of *Shigella* infection, loss of TLR signaling by hematopoietic cells also resulted in elevated *Shigella* IEC CFU (Fig. 5B) and enhanced pathology (Fig. 5C and extended data Fig. 5A).

### IL-12 induces IFN-γ production

IFN-γ can be induced by different cytokines, including IL-12, IL-15, IL-18, and type I IFN^49,50^. IL-18 neutralizing antibody administered intraperitoneally during *Shigella* infection of *Nlrc4*^−/−^ *Casp11*^−/−^ mice had no effect on IEC CFU burdens or pathology (Fig. S5B). Conversely, administration of IL-12 neutralizing antibody significantly increased the susceptibility of *Nlrc4*^−/−^ *Casp11*^−/−^ mice to *Shigella* infection (Fig. 5D). Mice treated with anti-IL-12 showed considerably higher intraepithelial bacterial counts compared to those given an isotype control (Fig. 5D). Furthermore, neutralizing IL-12 exacerbated weight loss and aggravated inflammatory symptoms of shigellosis, including cytokine levels and markers such as MPO and lipocalin (Fig. S5E). Notably, IL-12 neutralization resulted in a significant reduction of IFN-γ in the LAP (Fig. 5E) and the exacerbation of bacillary dysentery observed in these mice mirrors the disease elevation observed in *Ifngr1*^−/−^*Nlrc4*^−/−^*Casp11*^−/−^ mice. Moreover, treatment with anti-IL-12 had no effect on the severity of *Shigella* infection in the absence of IFN-γR (Fig. 5F), consistent with a model in which the main role of IL-12 in mediating resistance to *Shigella* is via induction of IFN-γ.

**Figure S5:**
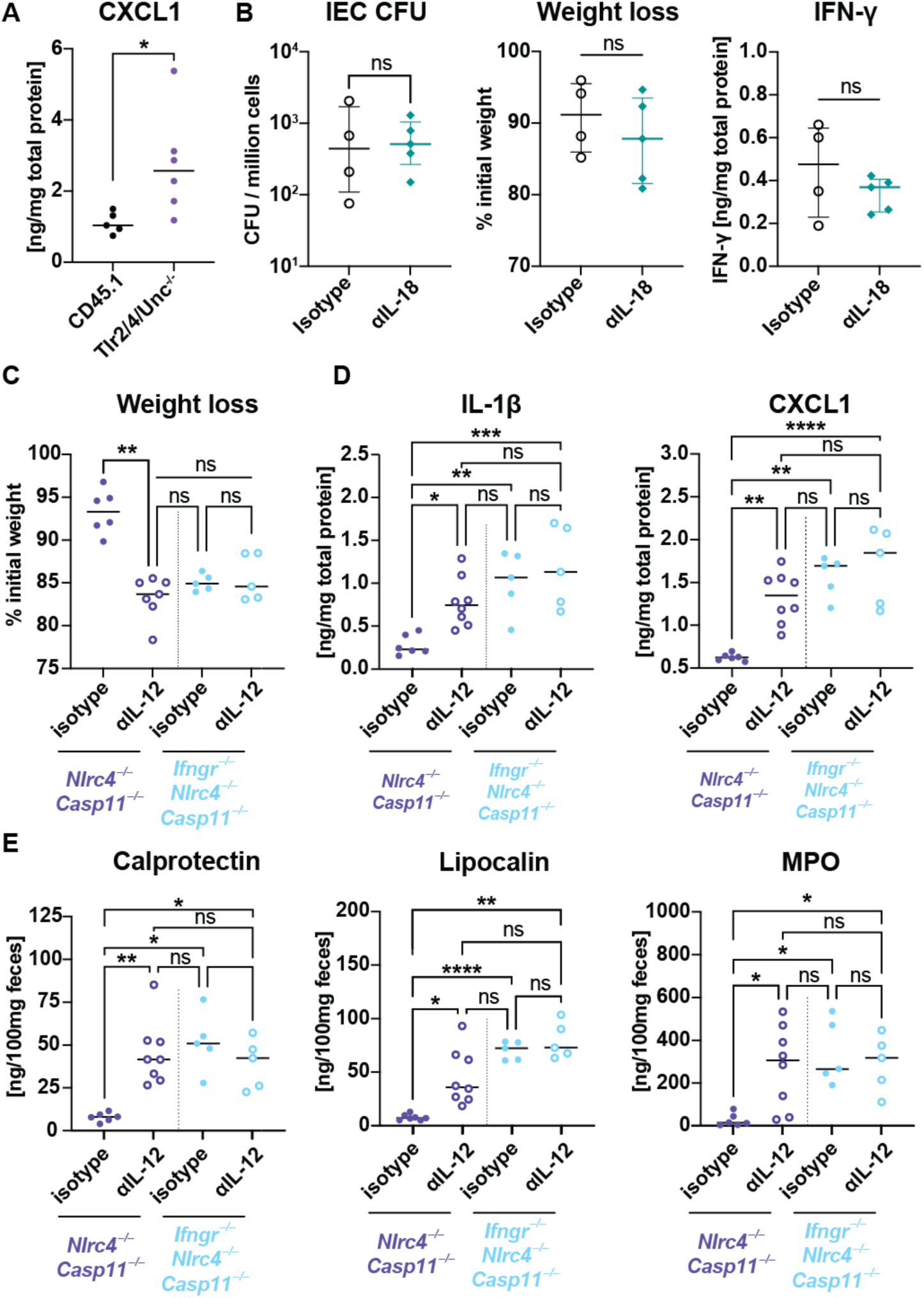
IL-12 neutralization phenocopies IFN-γR deficiency, whereas IL-18 neutralization has no effect. **(A)** ELISA for CXCL1 in the LAP 48 hours post-infection in irradiated *Nlrc4*^−/−^*Casp11*^−/−^, receiving *Tlr2/*4*-Unc*^−/−^ or CD45.1-WT bone marrow. **(B)** IEC CFU, weight loss 48 hours after infection, IFN-γ levels in the LAP observed in *Nlrc4*^−/−^*Casp11*^−/−^ mice treated with an IL-18 neutralizing antibody or an isotype control. **(C)** Animal weight 48 hours after infection normalized to initial weight, **(D)** IL-1β and CXCL1 levels in LAP, **(E)** Calprotectin, lipocalin and MPO in the feces of infected *Nlrc4*^−/−^*Casp11*^−/−^ or *Ifngr^−/−^Nlrc4*^−/−^*Casp11*^−/−^ mice additionally treated with αIL-12 or isotype control. ns=not significant, *p<0.05, **p<0.01, ***p<0.001, ****p<0.0001 according to t-test (B), ANOVA with Tukey’s multiple comparisons (C,B,E). n=5 (A), n=4-5 (B), n=5-8 (C,D,E).

## Discussion

Bacillary dysentery is a self-limiting disease in most healthy adults. However, the lack of suitable animal models has hindered our ability to determine the key processes that control *Shigella* replication and disease symptoms *in vivo*. We previously showed that *Shigella* can successfully invade and replicate in epithelial cells of mice lacking the NLRC4 inflammasome^14,15^. Mice deficient in both NLRC4 and Caspase-11 are even more susceptible to *Shigella*, and exhibit all pathological aspects of human shigellosis^15^. Here, we show that infected *Nlrc4^−/−^Casp11^−/−^* mice recover from dysentery within five days, similar to the typical disease course in humans (Fig. 1A, B).

The swift recovery of infected mice within a few days indicates that the processes that ultimately curtail *Shigella* growth must already be activated in the early stages of the disease, consistent with a pivotal role of the innate immune response. Furthermore, as epithelial cells of the large intestine are the predominant niche for *Shigella*, these innate responses must ultimately restrict or prevent intracellular bacterial replication in IECs. The top differentially expressed genes in IECs after *in vivo* infection are characteristic of an IFN-γ response (Fig. 1C, D, S1B). Indeed, we were able to show that mice failing to mount a response to IFN-γ were incapable of recovering from a *Shigella* infection (Fig. 3A, B). Infection of mice treated with an IFN-γ-neutralizing antibody, or mice deficient of the IFN-γ receptor, were highly susceptible to shigellosis, presenting with severe diarrhea, exacerbated inflammation and 1,000 times higher CFUs within IECs compared to infected control animals (Fig. 2B-I, S2A, S3A).

Using reciprocal bone marrow chimeras, we found that IFN-γ exerts its effects directly on non-hematopoietic cells (Fig. 3D). Given that *Shigella* replicates within IECs, the most parsimonious explanation for our data is that IFN-γ is acting directly on infected IECs. Furthermore, this hypothesis is underscored by the restrictive effect of IFN-γ pretreatment on the intracellular replication of *Shigella* in human colonic organoids (Fig. S1D). However, more indirect mechanisms for the action of IFN-γ are also possible. Ultimately, specific deletion of *Ifngr1* from IECs (on an *Nlrc4*^−/−^*Casp11*^−/−^ background) will be necessary to confirm whether IFN-γ signaling in IECs is necessary for *Shigella* control *in vivo*.

IFN-γ has been identified as a key factor in restricting several intracellular pathogens, including *Mycobacterium tuberculosis*, *Listeria monocytogenes*, and *Legionella pneumophila*^51–57^. IFN-γ induces the expression of hundreds of interferon-stimulated genes (ISGs) to promote pathogen clearance. Several ISGs have been shown to confer antibacterial functions during *Shigella* infection *in vitro*^58–62^. The most prominent example is the family of guanylate-binding proteins (GBPs). GBPs bind and liberate LPS to promote CASP11- and CASP4-mediated pyroptotic cell death^60,63–66^. In addition, the binding of GBPs to LPS forms a cage-like structure, trapping intracellular bacteria and inhibiting actin-dependent motility^67,68^. However, *Shigella* disrupts the GBP coat by secretion of an E3 ubiquitin ligase effector protein, IpaH9.8, which ubiquitylates GBPs and targets them for proteasome-dependent degradation^62,65,68^. Additionally, the *Shigella* effector OspC3 blocks Caspase-4/11-mediated inflammatory cell death^21,31,32^. Other ISGs, such as Viperin, Apolipoprotein 3 (APOL3), and the E3 ligase RNF213, have been attributed to restricting intracellular *Shigella* to some degree^58,59,69,70^. However, *Shigella* expresses a range of effector proteins that counteract these activities. For instance, IpaH1.4 directly antagonizes RNF213 by mediating its proteasomal degradation^69,70^. Despite *Shigella* antagonism of ISGs, our study in mice identified IFN-γ responses as central in restricting *Shigella in vivo* and in human colonic organoids. Further experiments will focus on determining which ISGs restrict *Shigella* and their specific mechanism.

Despite the widely held belief that neutrophil recruitment and transepithelial migration into the gut lumen are significant drivers of pathology during shigellosis^1,7^, we find no major role for neutrophils during *Shigella* infection (Fig. 4C, D, F, Fig. S4D, E). Importantly, we observed extensive recruitment of neutrophils into the lamina propria (Fig. S4A), and the presence of the neutrophil-specific enzyme myeloperoxidase in the feces. These observations suggest that neutrophils are recruited to the intestine in our model, and cross the epithelial layer, as in humans, in which the presence of neutrophils in the stool is a clinical marker for shigellosis^1,71,72^. Nonetheless, the only noticeable effects we observed upon neutrophil depletion were the absence of pus (Fig. S4F) and undetectable MPO levels in the gut lumen, without any impact on the weight loss or the bowel inflammation markers lipocalin and calprotectin (Fig. 4C, D, F and S4D, E). Importantly, neutrophil depletion during infection did not affect IEC bacterial counts. Our results underline the importance of using genetic models to distinguish causal versus correlational roles of host factors during infection.

It is intriguing that neutrophils appear neither to exacerbate disease nor restrict the growth of *Shigella*. While numerous studies highlight the critical role of neutrophils in eliminating invading microbes and resolving inflammation (as reviewed by Fournier *et al.*^73^), in the case of *Shigella* infection, neutrophils seem to have negligible impact on bacterial restriction. This might be explained by the ability of *Shigella* to effectively induce cell death of phagocytic cells^7,71^ or by *Shigella* evasion of phagocytes by replication in epithelial cells. Alternatively, neutrophils may enhance bacterial clearance, but this anti-bacterial function is offset by a distinct pro-bacterial (e.g., immunoregulatory) effect of neutrophils, such that there is no net change in bacterial burdens in neutrophil-depleted mice. It may also be the case that neutrophils play important roles in preventing systemic spread of *Shigella*, or in the late stages of the infection, neither of which were examined in our study.

*Shigella* rapidly induces macrophage cell death at early stages of infection^43,74,75^. It is generally accepted that *Shigella* induces macrophage death to enable the subsequent infection of epithelial cells^7,42^. Rapid induction of macrophage death is also believed to limit the ability of macrophages to restrict bacterial replication. However, a recent study using a zebrafish model of *Shigella* infection found a host-protective role for macrophages^76^. Thus, it remains unclear whether *Shigella*-mediated macrophage cell death is beneficial to the pathogen or to the host and whether macrophages participate in host defense against *Shigella* intestinal infections. In experiments with mice lacking monocytes and macrophages, we detected increased disease severity and colonization of the intestinal epithelium (Fig. 4A, B, E, and S4C). We also noted similar pathology in mice deficient in TLR signaling (Fig. 5B, C). TLRs sense extracellular-derived microbial ligands, and since *Shigella* is a pathogen that replicates in the cytosol, it is generally believed that cytosolic sensing pathways (e.g., NOD1/2^77^and ALPK1^78^) are more important in infected cells than TLRs in initiating innate immunity to *Shigella*. However, it is likely that microbial ligands are released extracellularly during infection, and thus, bystander (uninfected) macrophages may sense and respond to *Shigella*. We therefore propose a model in which the release of bacterial products from infected, pyroptotic macrophages facilitates the release of protective cytokines from noninfected bystander cells—most likely macrophages—in a TLR-dependent manner. Together, our data suggest a beneficial role for macrophages in host defense during *Shigella* infection.

Finally, our results demonstrate that macrophage activation is pivotal in the induction of adequate IFN-γ to restrict *Shigella* (Fig. 5A). Although some reports suggest that macrophages can directly produce IFN-γ^79,80^, our experiment using IL-12 neutralizing antibodies suggests that the secretion of IFN-γ is indirect and reliant on IL-12, a cytokine known to be induced in macrophages downstream of TLR activation. Further experiments will focus on the identification of the cell type producing and releasing IFN-γ in response to IL-12 during the *Shigella* infection *in vivo*.

In sum, our results necessitate revisions to the generally accepted model of *Shigella* pathogenesis. In particular, our results identify a critical function for macrophages, but not neutrophils, in host defense against *Shigella*. Given that infected macrophages are rapidly killed by *Shigella*, we propose that bystander (uninfected) macrophages are vital in orchestrating anti-*Shigella* innate immunity. Instead of directly killing *Shigella* by phagocytosis, we propose that a TLR–IL-12 circuit induces the expression of IFN-γ, which then acts on IECs to restrict *Shigella* replication. Ultimately, understanding how host immunity coordinates clearance of *Shigella* may be essential for the development of an effective *Shigella* vaccine.

## Material and Methods

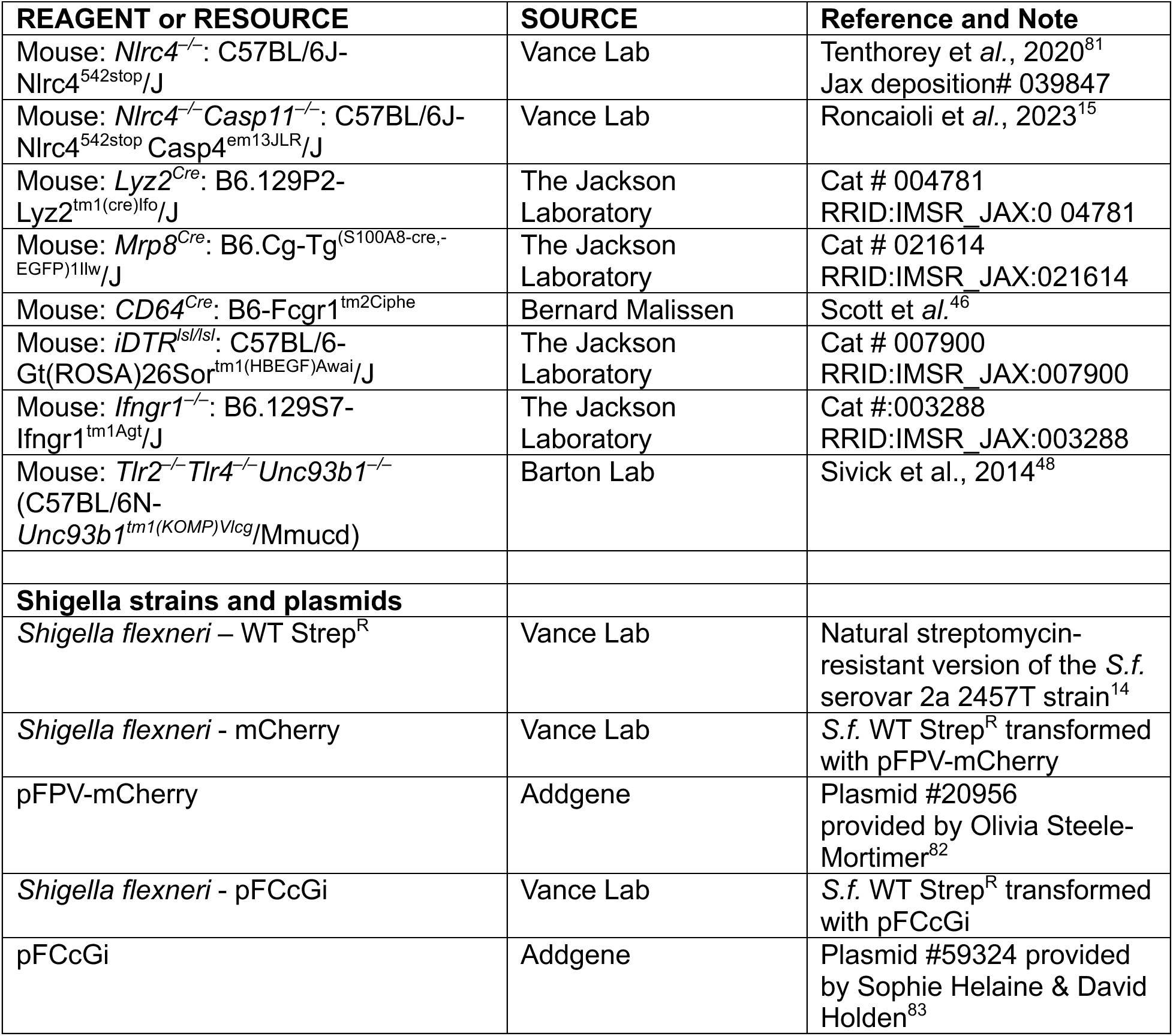

### Animals

Mice were maintained under specific pathogen-free conditions and housed with a 12-hour light-dark cycle and standard chow diet (Harlan irradiated laboratory animal diet) ad libitum in accordance with the regulatory standards of the University of California Berkeley Institutional Animal Care and Use Committee. All mice were sex- and age-matched and were 6-12 weeks old when infected (except mice receiving bone marrow transplantation, see details below). Both male and female mice were used in all experiments. Littermate controls were used or, if not possible, mice were co-housed for at least three weeks prior to infection.

C57BL/6J-*Nlrc4*^542stop^*Casp4*^em13JLR^/J (*Nlrc4^−/−^Casp11^−/−^*) animals were previously generated and described by Roncaioli et *al*.^15^ via targeted CRISPR-Cas9 mutagenesis of *Casp11* in existing *Nlrc4^−/−^*(C57BL/6J-Nlrc4^542stop^/J)^81^ mice.129P2-*Lyz2*^tm1(cre)Ifo^/J (*Lyz2^Cre/+^*)^84^, B6.Cg-Tg^(S100A8-cre,-EGFP)1Ilw^/J (Mrp8*^Cre^*)^45^, C57BL/6-Gt(ROSA)26Sor^tm1(HBEGF)Awai^/J (iDTR*^lsl/lsl^*_)_^85^, B6.129S7-*Ifngr1*^tm1Agt^/J (*Ifngr1^−/−^*)^86^ mice were purchased from Jackson Laboratories. The *Ifngr1^−/−^Nlrc4^−/−^ Casp11^−/−^*mouse line was generated by mating *Ifngr1^−/−^* and *Nlrc4^−/−^Casp11^−/−^*mice. B6-*Fcgr1*^tm2Ciphe^ (*CD64^Cre/+^*)^46^ mice were generated by Bernard Malissen at Centre d’Immunologie de Marseille-Luminy and provided by Yasmine Belkaid at the National Institutes of Health. *iDTR^lsl/lsl^Nlrc4^−/−^Casp11^−/−^* were generated by crossing *iDTR^lsl/lsl^* mice, obtained from the Jackson Laboratory, with our *Nlrc4^−/−^Casp11^−/−^*mice. Subsequently, *Lyz2^Cre/+^iDTR^lsl/lsl^Nlrc4^−/−^Casp11^−/−^*, *CD64^Cre/+^iDTR^lsl/lsl^Nlrc4^−/−^Casp11^−/−^*and *Mrp8^Cre/+^ iDTR^lsl/lsl^ Nlrc4^−/−^Casp11^−/−^*lines were generated by crossing *iDTR^lsl/lsl^ Nlrc4^−/−^Casp11^−/−^* mice to animals form the *Lyz2^Cre/+^*, *CD64^Cre/+^* or *Mrp8^Cre/+^* line, respectively. C57BL/6N-*Unc93b1^tm1(KOMP)Vlcg^*/Mmucd (*Unc93b1*^−/−^) mice were obtained from the Mutant Mouse Resource and Research Center (MMRRC) at the University of California, Davis, and originally donated to the MMRRC by David Valenzuela of Regeneron Pharmaceuticals^87^ and after crossing to *Tlr2^−/−^Tlr4^−/−^* mice, *Tlr2^−/−^Tlr4^−/−^Unc^−/−^* animals were generously provided by Gregory Barton at the University of California, Berkeley.

### *Shigella* cultivation and preparation for *in vivo* infections

If not stated otherwise, infection experiments were conducted with a natural streptomycin-resistant strain of *Shigella flexneri* serovar 2a 2457T^32^ (*S.f.*). *S.f.* was grown on tryptic soy broth (TSB, BD Bacto # DF0370-07-5) agar plates containing 0.01% congo red (CR, Sigma-Aldrich # C6767) and 100 µg/mL streptomycin sulfate at 37°C. For infections, a single CR-positive colony was picked from a streak not older than a week, inoculated into 5 mL TSB supplemented with 100 µg/mL streptomycin, and incubated overnight under constant agitation at 37°C. 16 hours later, the culture was back-diluted 1:100 in 5 mL fresh TSB + 100 µg/mL streptomycin and incubated for another ∼3 hours under constant shaking at 37°C. Upon reaching an OD_600_ of 1, bacteria were pelleted by centrifugation with 3,000×*g* for 8 minutes, washed twice with PBS, and resuspended in the same volume of pharmaceutical grade PBS (USP) for infection by oral gavage. From several experiments that tracked the correlation between OD_600_ and colony-forming units (CFU), we can estimate that an OD_600_ equals roughly an infection dose of 1-2×10^8^ CFU/mL. Nevertheless, the actual infectious dose was determined for each experiment by serial dilution and plating on TSB agar plates containing 0.01% CR.

### *In vivo* infection

One day before infection, littermates or cohoused mice were deprived of food and water in the morning and 4-6 hours later orally gavaged with 100 µL of 250 mg/mL streptomycin sulfate dissolved in pharmaceutical grade PBS, after which mice were again given food and water. On the day of infection, food and water were removed again from the cage, and 4-6 hours later, mice were orally gavaged with 100 µL of 10^8^ CFU/mL *S.f.,* prepared as described above. In compliance with our Animal Utilization Protocols, infected mice were visually examined twice per day and their change in weight was monitored every 24 hours. A loss of more than 25% of the initial weight measured on the day of infection (day 0), or signs of severe sickness such as pronounced hunching, combined with shivering or breathing distress, were considered humane endpoints at which mice were euthanized.

For *in vivo* antibody-mediated cytokine neutralization, 500 μg of anti-IFN-γ antibody (clone XMG1.2, Bio X Cell #BE055), anti-IL12 (clone C17.8, Bio X Cell #BE0051), anti-IL-18 (clone YIGIF74-1G7, Bio X Cell #BE0237),) and polyclonal Armenian hamster IgG isotype control antibody (clone LTF-2, Bio X Cell #BE090) were administered by daily intraperitoneal injection starting on the day of infection. For antibody-mediated depletion of neutrophils, 500µg of anti-Ly6G (clone 1A8, Bio X Cell #BE0075-1) was injected daily by intraperitoneal injection starting one day before infection (at the time of streptomycin administration).

In experiments involving diphtheria toxin-mediated cell depletion, mice were injected i.p. with 30ng/g of DT (Sigma #D0564) or an equal volume of pharmacological PBS daily, starting one day before infection.

### Assessment of inflammation

If not indicated otherwise, 48 hours after infection, mice were sacrificed, their colon and cecum were isolated, and their lengths recorded. Subsequently, the tissue was cut longitudinally, and the most distal fecal matter from the colon was collected, some of which was spread on detection tabs from a Hemoccult blood testing kit (Pro Advantage #P080018). The rest was then transferred into a pre-weighed 2 mL tube. The fecal matter was homogenized in 1 mL of PBS containing protease inhibitors (Roche #04693159001) using a polytron homogenizer. To determine the fecal CFU, serial dilutions were made in PBS and plated on TSB containing 0.01% CR and 100 µg/mL streptomycin sulfate. For lipocalin, calprotectin, and MPO ELISAs, samples were centrifuged at 13,000×*g* for 5 minutes and supernatants were analyzed in duplicates with R&D sandwich ELISA kits according to the manufacturer protocol.

A four-step scale was applied to quantify the presence of occult blood. A value of 0 indicates that the Hemoccult blood test was negative, 1 indicates a faint blue staining, and 2 indicates an intense staining. Mice who experienced severe hemorrhagic conditions with clearly visible, macroscopic blood in the lumen of the colon were assigned a score value of 3. Assessments were made on blinded samples.

### IEC CFU

To determine the levels of intra-epithelial bacteria, the colon and cecum were isolated from infected mice 48 hours after infection as described above. After taking the fecal sample, the tissue was cleared of any remaining luminal content by washing in PBS. The tissue was collected in 5 mL RPMI (Thermo Fisher #21870-76) with 5% FBS, 2mM GlutaMax (Thermo Fisher #35050-61), 25mM HEPES (Thermo Fisher #15630-080), and 400 µg/mL gentamicin (Thermo Fisher #15710-064), and incubated for 1-2 hours at 4°C. Subsequently, tissue was washed six times in PBS, minced into approximately 1 cm pieces, and placed in 12 mL IEC stripping solution (HBSS, 25 HEPES, 2mM Glutamax, 50µg/mL Gentamicin, 2mM DTT, and 5mM EDTA) within a 50 mL Erlenmeyer flask and incubated for 30 minutes at 37°C and constant stirring at low speed (220rpm). The supernatant was passed through a 100 µm cell strainer, and the retained tissue was transferred back into the flask and mixed with 10 mL ice-cold PBS. The flask was sealed with a rubber stopping, shaken vigorously for 20 seconds, and combined with the previous extraction by passing through the same cell strainer. The remaining tissue was saved for further processing (see LAP ELISA or Flow staining). Next, the obtained IEC fraction was incubated with 50 µg/mL gentamicin on ice for 20 minutes, centrifuged 500×*g* for 5 minutes at 4°C, and washed twice with ice-cold PBS. Before the last centrifugation, a small aliquot of cells was taken for cell counting. Finally, the pellet was resuspended in 1 mL of 1% TritonX-100 (Fisher #BP151-100) and IEC-CFU was determined by plating serial dilutions on TSB agar plates with 0.01% CR and 100 µg/mL streptomycin sulfate.

### Tissue ELISA

After extracting IEC cells, the remaining tissue was transferred into 14 ml round-bottom tubes containing 1 mL PBS with proteinase inhibitors (Roche #04693159001). After homogenization with a polytron homogenizer at 20,000 rpm, the suspension was centrifuged at 13,000×*g* for 5 minutes and the supernatant was analyzed in duplicates with R&D sandwich ELISA kits following the manufacturer protocol and normalized to the total protein concentration determined with the Pierce BCA protein assay kit (Thermo scientific #23225) according to the manufacturer protocol.

### CFU Spleen/MLN

The spleen and mesenteric lymph nodes were isolated from infected mice 48 hours after infection. Collected tissue was placed in RPMI with 5% FBS, 2mM GlutaMax, 25mM HEPES, and 400 µg/mL gentamicin for 1h. After washing 5× with PBS, tissue was homogenized using a polytron homogenizer and serial dilutions plated on TSB agar plates with 0.01% CR and 100 µg/mL streptomycin sulfate.

### IEC bead enrichment and RNAseq

The IEC faction was obtained as described above and further digested with Dispase-II to generate a single-cell suspension, as described by Gracz *et al*^88^. In brief, the pelleted IEC fraction was washed with 10 mL PBS with 10% PBS and resuspended in 10 mL of pre-warmed (37°C) HBSS containing 8 mg Dispase-II (Sigma #D4693). Tubes were incubated in a 37°C water bath for 10 minutes with vigorous shaking every 2 minutes. Next, 1 mL FBS with 500 µg DNase (Roche #11284932001) was added and incubated 3 minutes on ice. After washing with RPMI containing 10% FBS, cells were passed through a 40 µm cell strainer. Epithelial cells were enriched using the MojoSort™ Mouse CD326 (Ep-CAM) Selection Kit (BioLegend Cat #480141) following the manufacturer’s protocol for Positive Selection. Retained CD326^+^ cells were washed with 4mL RPMI containing 10% FBS and centrifuged with 1,000×*g* for 5 minutes at 4°C and lysed in 250 µL Trizol (Thermo Fisher #10296028). Samples were topped up with 100 µL DNase/RNase-free water and mixed with 200 µL Chloroform (Fisher Scientific #C298-500). The aqueous phase and mixed with an equal volume of ethanol. Subsequently, the RNA was isolated using the Monarch Total RNA Miniprep Kit (NEB #T2010S) following the manufacturer’s protocol with on-column DNase treatment. Obtained RNA was sent to Azenta for quality assessment, automated PolyA selection, library preparation, multiplexing and paired-end sequencing with a read length of 150 bases. After demultiplexing, raw data were pre-processed (including quality control as well as barcode, adaptor, and quality trimming) using FastQC (www.bioinformatics.babraham.ac.uk/projects/fastqc/) and cutadapt, mapped to the Mus_musculus.GRCm38.96 genome assembly using Kalisto (0.44.0 via Bioconda). Overall, each sample contained between 24 million and 32 million reads. Differential expression analysis was performed using DESeq2 (V1.44.0)^89^ with lfc shrinkage correction in R project version 4.4.1 with RStudio version 2024.04.2+764 (Foundation for Statistical Computing, Vienna, Austria, www.R-project.org/). For gene ontology (GO) enrichment analysis genes with differential expression values padj < 0.001 and log_2_ fold change > 1 were selected and enrichment analysis was performed using gseGO of the clusterProfiler (4.12.2) package for R with a gene set size <150.

### LAP flow

To prepare cells from the lamina propria for flow cytometry, the colon and cecum were isolated from infected mice and striped from IEC as described above. The collected remaining tissue was minced thoroughly with scissors and transferred into 50 mL Erlenmeyer flasks. After adding 10 mL of pre-warmed HBSS with 100 µg/mL Liberase TM (Sigma #5401127001) and 5 µg/mL DNase, tissue was digested for 45 minutes under constant stirring at 37°C. The cell suspension was passed through a 70 µm strainer and washed twice with cold RPMI containing 10% FBS. Epithelial and dead cells were removed using a 70% and 40% two-phase Percoll gradient. Cells from the interface were subsequently collected, washed with PBS and stained with Ghost Dye™ Red 780 (Tonbo #SKU 13-0865-T100) before blocking the FcγII/III receptor with an anti-CD16/anti-CD32 antibody (BioLegend, #156604). Samples were stained for 45 minutes to an hour at room temperature in FACS buffer (PBS with 5g/L BSA and 2g/L sodium azid) with the following antibodies: BV785-labeled CD45 (Clone 104, BioLegend #109839), BUV496-labeled CD3 (Clone 145-2C11, BD #612955), APC-eFlour-labeled B220 (Clone RA3-6B2, Thermo Fisher #47-0452-82), PE-labeled CD11b (Clone M1/70 Thermo Fisher #12-0112-82), FITC-labeled Ly6G (Clone 1A8, BD #551460), APC-labeled Ly6C (Clone HK1.4, BioLegend #128016), MHCII (Colon M5/114.15.2, BioLegend #107626), BUV605-labeled CD64 (Clone H1.2F3, BioLegend #104530), BV421-labeled F4/80 (Clone BM8, BioLegend #123137) and APC-Cy7-labeled EpCAM (Clone G8.8, BioLegend #118218). Stained samples were washed twice and fixed with cytofix/cytoperm (BD biosciences #554722) for 10 minutes at room temperature before measured on an Aurora (Cytek) flow cytometer. Data were analyzed with Flowjo version 10 (BD Biosciences).

### Blood Staining

Terminal blood sampling was performed by retro-orbital sinus puncture of anesthetized mice with a heparinized capillary (#22-260950 Fisher scientific). 15µL of the collected blood was immediately mixed with 100µL PBS containing 100U/mL heparin (#H0878-100KU Sigma) and 25µL counting beads (AccuCheck Counting Beads, Thermo Fisher #PCB100). After centrifugation and washing with FACS buffer, cell pellet was stained with 25µL of antibody mix (APC-labeled Ly6C (Clone HK1.4, BioLegend), FITC-labeled Ly6G (Clone 1A8, BD #551460), PE-labeled CD3 (ebio #12-0031-81 Clone 14-2011)) along with the viability dye (Tonbo #13-0658-T) and FcγII/III-block. Subsequently, cells were fixed cytofix/cytoperm (BD biosciences #554722) for 10 min and red blood cells were lysed with ACK buffer (Thermo fisher #A1049201). After washing twice with FACS buffer, samples were analyzed with an Aurora (Cytek) or a BD LSR Fortessa flow cytometer.

### CT26 infection

CT26 were purchased from the Berkeley Cell Culture Facility and cultured in RPMI with 10% FBS, 2 mM Glutamax (Thermo Fisher #35050061), 10 mM HEPES (Thermo Fisher # 15630106), 1 mM sodium pyruvate (Thermo Fisher #11360070) and Penicillin-Streptomycin (Thermo Fisher #15070063). 10^6^ CT26 cells were seeded into one well of a 6-well tissue culture treated plate in media without Penicillin-Streptomycin one day before the infection. *Shigella flexneri* -pFCcGi (constitutively expressing mCherry and an arabinose inducible GFP) was grown as described above. CT26 cells were pre-treated with 10 ng/mI IFN-γ (Peprotech #315-05-100UG) or left untreated for 16 hours and were spin-infected (600xg 10min at 37°C) with mid-log phase *Shigella* at an MOI of 1. After 45 minutes, cells were washed twice with PBS and medium exchanged to CT26 medium supplemented with 100 µg/ml gentamicin (Thermo fisher # 15710064) and 0.4% L-Arabinose (Sigma #A3256-25G). After 3 hours cells were isolated by trypsinization, washed, passed through a 70µm cell strainer, stained with Ghost Dye™ Red 780 (Tonbo #SKU 13-0865-T100), fixed with cytofix/cytoperm (BD biosciences #554722) for 10 min and analyzed with a BD LSR Fortessa flow cytometer.

### Human organoid maintenance and infection

Deidentified human colonic organoids were a gift from Scott B. Snapper, M.D., Ph.D. and established from a rectal biopsy obtained during a routine diagnostic endoscopy in a pediatric subject under Boston Children’s Hospital IRB protocol and cultured with methods modified from Sato *et al*.^90^. Briefly, organoids were maintained in 50μL Matrigel domes with human IntestiCult™ Organoid Growth Medium (STEMCELL #6010) supplemented with 10 µM of the ROCK1/ROCK2 inhibitor Y-27632 and Penicillin-Streptomycin. 16 hours before infection, organoids were stimulated with 10 ng/ml IFN-γ (Peprotech # 300-02-100UG). Before infection domes were washed with PBS and disrupted with a pipette tip and the Matrigel was digested with 0.25% Trypsin at 37°C for 5min. After washing with media, organoids were infected with mid-log phase *Shigella flexneri* -pFCcGi at an MOI of approximately 1. After 45 minutes, organoids were washed twice with PBS and medium exchanged to IntestiCult medium supplemented with 100 µg/ml gentamicin (Thermo fisher # 15710064) and 0.4% L-Arabinose (Sigma #A3256-25G). After 3 hours organoids were washed and, stained with Ghost Dye™ Red 780 (Tonbo #SKU 13-0865-T100). After single-cell digestion with TrypeLE (Thermo fisher #12604013) at 37°C for 5 minutes, cells were fixed with cytofix/cytoperm (BD biosciences #554722) for 10 min, passed through a cell strainer and analyzed with a BD LSR Fortessa flow cytometer.

## Acknowledgment

We thank Scott B. Snapper, David Breault, Daniel Zeve, and Richelle Bearup and the Clinical Translational Research Program and the Organoid Core of the Harvard Digestive Disease Center (supported by NIH grant P30DK034854), Division of Gastroenterology, Hepatology, & Nutrition, Boston Children’s Hospital for providing deidentified human colonic organoids.

## Competing interests

Russell E Vance is an SAB member of X-biotix and Tempest Therapeutics. The other authors declare that no competing interests exist.

## Funding

R.E.V. and G.M.B. acknowledge research support from Investigator Awards and an Emerging Pathogens Initiative Award from the Howard Hughes Medical Institute. R.E.V. is also supported by NIH grants AI075039 and AI155634. G.M.B. acknowledges research support from NIH/NIAID grant R01AI072429 and C.F.L. acknowledges research support from NIH grant AI169795.

